# PDE4 Inhibitor Apremilast Rebalances Inflammatory Responses to Pseudomonas aeruginosa Infection in CF Rats

**DOI:** 10.64898/2026.02.08.704692

**Authors:** Linto Antony, Lawrence Rasmussen, Denise Stanford, Antonio Allen, Daniel Kennedy, Levi W. Brewer, Aditi Shanbhag, Jennifer LaFontaine, S. Vamsee Raju

## Abstract

CFTR modulator therapies have transformed CF care, yet chronic airway inflammation persists in many people with cystic fibrosis (pwCF) even after long-term highly effective modulator therapy (HEMT). Because of the adverse side effects or the incompatibility with CFTR modulators, the use of traditional anti-inflammatory therapies is very limited in CF. Hence, new therapeutic strategies that rebalance inflammation without worsening infection with immunosuppression are needed. We evaluated the selective phosphodiesterase 4 (PDE4) inhibitor apremilast (Apr) for its ability to modulate dysregulated inflammation in humanized CF (G551D) rats acutely challenged with *Pseudomonas aeruginosa*. Apr is an approved anti-inflammatory therapeutic strategy for several chronic inflammatory conditions, but it has not been well studied in CF. In the humanized CF (G551D) rats, a short prophylactic Apr regimen significantly preserved lung function and reduced lung injury, accompanied by broad modulation of inflammatory responses, notably within Th1 and Th17 axes. Importantly, Apr did not cause a significant increase in bacterial burden. Just as importantly, Apr did not reduce CFTR mRNA or protein in vivo, and it increased G551D-CFTR phosphorylation critical for channel gating in vitro, supporting mechanistic compatibility with HEMT. These findings suggest Apr as a potential adjunct to CFTR modulators to rebalance airway inflammation while preserving host defense.

## Introduction

Altered airway physiology, caused by mutations in the cystic fibrosis transmembrane conductance regulator (CFTR) gene and the corresponding dysfunctional chloride ion transport by defective CFTR protein, predisposes people with cystic fibrosis (pwCF) to inflammation and recurrent infections^1, 2^. Persistent inflammation, triggered by hypoxic stress due to hyperviscous and obstructive mucus in the airway, even in the absence of an active infection, is a hallmark feature of cystic fibrosis (CF)^2–4^. Progressive decline in the pulmonary functions attributed to either chronic inflammation and/or infection represents the chief determinants of morbidity and mortality in pwCF^5, 6^. Inflammation, along with airway obstruction and infection, forms a triad of pathophysiological factors around the central genetic cause of CF. Since these factors are interconnected, managing one factor affects the sequelae of the other component^7^. Given these constraints, adjunct anti-inflammatory strategies that are mechanistically compatible with CFTR modulators and avoid overt immunosuppression are needed.

Modulator therapies targeting the CFTR protein were a turning point in CF pharmacotherapy that fundamentally transformed the landscape of CF disease management. After its approval for CF treatment, highly effective modulator therapy (HEMT) made substantial improvements in pulmonary function, median survival age, and a decline in pulmonary exacerbations (PEx) and lung transplantation rates over the past few years, as documented in the 2024 Patient Registry Highlights by the Cystic Fibrosis Foundation^8^(https://www.cff.org/medical-professionals/2024-patient-registry-highlights). Despite the transformational improvements in the prognosis of pwCF and the evolvement of the CF care model, modulator therapies were incapable of restoring lung function to normative levels and failed to eradicate CF-associated pathogens and inflammation from CF lung^9–12^. Although significant reductions in inflammation and improved clinical outcomes were achieved, persistent infection and inflammation remain as enduring challenges even after long-term modulator therapy ^13–17^. This inefficiency of modulators was also shown in pre-clinical studies, where treatment with the CFTR-potentiator ivacaftor (Iva) could not resolve the inflammation triggered by LPS^18^. Furthermore, Iva treatment altered the phenotype of *Staphylococcus aure*us (*S. aureus*) to a more persistent one rather than decreasing its burden in the CF lung^19^. Corroborating with these evidences, a recent report from the PROMISE study demonstrates that even after long-term therapy with triple combination (ETI) HEMT, inflammation persists in the CF lung due to non-clearance of *Pseudomonas aeruginosa (P. aeruginosa)* ^20^. Hence, comprehensive therapeutic strategies that help to clear both inflammation and infection in the CF lungs are utmost warranted in CF disease management.

Intensive antimicrobial therapies are effective but provide only a transient pathogen clearance because of the post-antibiotic resilience exhibited by major CF pathogens such as *P. aeruginosa* and *S. aureus*. The adaptation of these pathogens, along with the acquisition of antibiotic resistance and the development of immune evasion, as well as negative effects on microbial diversity in the respiratory system, are other disadvantages of antibiotic treatment^21, 22^. Clinically, CF patients harboring resistant pathogenic strains tend to present with severe baseline pulmonary disease, experience accelerated decline in lung function and have a higher propensity for lung transplantation^23^. Studies suggest that a long-term antibiotic regimen can protect the CF lungs from infection but fails to prevent the progression of inflammatory lung pathology^4^ and does not improve lung function during exacerbations^24^. While the emergence of antimicrobial resistance remains the foremost concern for antibiotics, the pharmacological incompatibility between these agents and CFTR modulators constitutes an equally critical obstacle to effective combination therapy. Reports of adverse clinical outcomes from drug interaction when the CFTR modulator Ivacaftor was co-administered with a commonly used antimicrobial agent, rifampin, serve as the best example of this^25^. Therefore, anti-inflammatory therapy remains the most plausible comprehensive treatment option complementing CFTR modulators.

Persistent proinflammatory microenvironments in pwCF begin to develop early in life, potentially contributing to disease progression^26–29^. Therefore, there is a continual need for anti-inflammatory therapies in managing CF disease. Due to the adverse side effects and serious outcomes linked to excessive suppression of inflammation, anti-inflammatory therapeutic options available for the CF community are currently very limited^30–33^. Most of the approved drugs within that limited range belong to steroidal or NSAID therapies; even then, they have failed to achieve improved clinical outcomes in pwCF with PEx^34, 35^. Moreover, in vitro studies demonstrate that agents from these classes of anti-inflammatory drugs restrict the response of CF epithelial cells to CFTR modulators during inflammation^36^. Thus, finding an anti-inflammatory therapy better suited to CF is equally important as correcting the underlying genetic defect, especially when pwCF have to deal with prolonged inflammation during their extended lifespan.

In this study, we investigate the therapeutic potential of selective phosphodiesterase 4 (PDE4) inhibition in CF using apremilast (Apr), an FDA-approved anti-inflammatory agent for psoriasis and psoriatic arthritis. We hypothesized that Apr would attenuate inflammation and preserve lung function in CF without exacerbating pathogen burden. To test this, we administered a short prophylactic Apr regimen to CF rats expressing human G551D-CFTR before an acute *P. aeruginosa* challenge. This approach evaluates whether PDE4 inhibition can rebalance host responses in a manner compatible with CFTR modulation. This strategy holds promise for repurposing an established drug to address unmet needs in CF management, potentially offering a safe, cost-effective, and rapidly deployable therapeutic option.

## Materials and Methods

### Animal studies

To study the protective role of selective inhibition of PDE4 using apremilast (Apr)against infection in CF lung, we used both wild-type (WT) and transgenic Sprague-Dawley rats carrying the human CFTR gene with G551D mutation (hG551D CF rats, referred as CF rats from here onwards) obtained from the Cystic Fibrosis Research Center (CFRC) Core at UAB. Depending on the availability, both male and female rats within an age range of 16-20 weeks were included in the study and maintained on ad libitum food and water. CF rats were given additional dietary supplements such as DietGel 76A (ClearH2O, Westbrook, ME, USA) and Go-LYTLEY (Braintree Laboratories, Inc, Braintree, MA, USA) electrolyte supplements in water at concentrations of 25% and 50%, respectively, as previously described^37^ to prevent any loss of body condition and CF-related gut obstruction. All procedures involving animals were performed in accordance with the animal ethics and welfare policies approved by the Institutional Animal Care and Use Committee (IACUC) at UAB.

### Bacterial culture and inoculum preparation

An overnight culture of *Pseudomonas aeruginosa* (PAO1 strain, PerkinElmer) in Luria-Bertani (LB) broth (Difco, Miller) was plated on LB plates and incubated overnight at 37°C. The cells on the plate were then resuspended in sterile 1X PBS to achieve a final bacterial load of 10^7^ CFU/mL. For acute infection in CF rats, 200μL of this bacterial suspension was inoculated intratracheally under isoflurane anesthesia.

### Animal experiments

Following a seven-day acclimatization period, CF rats were randomly allocated into two groups: one received a vehicle (Veh) comprising 1% methyl cellulose in Carbowax, and the other received Apr at a dose of 10 mg/kg, both administered orally once daily for three consecutive days. One hour post-final administration, the rats were intratracheally inoculated with approximately 10^6^ CFU of *P. aeruginosa* (PAO1 strain, PerkinElmer) under isoflurane anesthesia. Twenty-four hours after infection, the animals were sedated with xylazine and ketamine and maintained on isoflurane for lung function assessment. Subsequently, rats were euthanized through cardiac puncture for sample collection. Untreated WT and CF rats served as sham controls and were used to enable comparisons of genotype and treatment effects wherever is needed.

### In vivo respiratory mechanics

We used flexiVent oscillometry (flexiVent FX; SCIREQ Inc., Montréal, Canada), following the manufacturer’s instructions with slight modifications. Animals were anesthetized via intraperitoneal injection of a xylazine–ketamine mixture (1mL/Kg) at doses of 2.5 mg/kg and 60 mg/kg, respectively. A tracheostomy was then performed, and the animals were intubated with 14-G tubes (SCIREQ, XS-IT14). Respiratory mechanics were assessed while ventilating the animals using a computer-controlled flexiVent system. Equipped with a customized module (FX4), the Flexivent system can intersperse mechanical ventilation with a variety of volume- and pressure-controlled maneuvers, assisted by a precision piston pump. Flexiware derives pulmonary functions in these intubated rats using the single-frequency forced oscillation technique (FOT).

### Bronchoalveolar lavage fluid (BALF) collection and analyses

BALF collection was performed on the right inferior lobe following injection of 4 mL of cold phosphate-buffered saline (PBS). To ensure isolation of the targeted lobe, the right superior and middle lobes, as well as the contralateral lung, were clamped prior to lavage. Post-lavage, aliquots were collected for total cell counting via an automated cell counter (TC20, BIO-RAD) and differential cell analysis by fixing cells on cytospin slides. The residual BALF was centrifuged at 1800 rpm for 5 minutes to pellet cells for flow cytometry analysis, while the supernatant was aliquoted and stored at -80°C for subsequent downstream assays.

The concentrations of key inflammatory cytokines in BALF were quantified utilizing a multiplexed bead-based assay on the Luminex 200 platform. Sample preparation was performed according to the manufacturer’s protocols using the MILLIPLEX MAP Rat Cytokine/Chemokine Magnetic Bead Kit (Millipore).

### Bacterial load enumeration

Undisturbed lung tissue portions, after weighing, were homogenized in 1X PBS using a bead beater. Serial dilutions of the tissue homogenate were then plated on *Pseudomonas* isolation agar (NutriSelect Plus, Millipore) and incubated for more than 18 hours at 37°C before enumeration.

### Lung tissue cell isolation for immunophenotyping

Fresh lung tissue specimens were initially rinsed in 1X PBS and minced into fine fragments using a sterile scalpel within a petri dish. The minced tissue was transferred to a 50 mL conical tube (Falcon) containing 1 mL of dissociation buffer composed of RPMI supplemented with 1.25 mg/mL collagenase and 150 U/mL DNase I (Sigma). The samples were incubated at 37°C for 20–30 minutes in a cell culture incubator with gentle agitation. To facilitate mechanical dissociation, undissociated tissue fragments were passed through a 70 μm cell strainer using a 20 mL syringe plunger. The resulting cell suspension was washed with FACS buffer (1X PBS, 2% FBS, 25 mM EDTA), followed by centrifugation at 1800 rpm for 5 minutes at 4°C. The cell pellet was resuspended in 5 mL of ACK lysis buffer and incubated at room temperature for 3-5 minutes to lyse red blood cells. The lysate was then diluted with three times its volume of FACS buffer and filtered through a 70 μm nylon mesh to eliminate residual tissue debris and clumps. The final suspension was centrifuged at 1800 rpm for 3 minutes at 4°C; the supernatant was discarded, and the pellet was washed repeatedly with FACS buffer to ensure cell purity. Cells were then resuspended in freezing media (90% FBS and 10% DMSO) and stored at -80°C until further processing.

### Flow cytometry

Cell staining and flow cytometry analyses were conducted as previously described^38^. Briefly, following Fc receptor blockade, cells were stained with a panel of innate immune cell surface markers by using antibodies to the following: CD45 (OX1, eBioscience), CD11b (OX-42, Thermo Fisher) CD43 (W3/13, BioLegend), CD161 (3.2.3, BioLegend) MHC II ((I-Ek, Miltenyi Biotec), and His48 (HIS48, eBioscience). Dead cells were stained by LIVE/DEAD™ Fixable Aqua Dead Cell Stain Kit (Invitrogen). Compensation controls were prepared using cells stained with single markers and unstained cells. After staining, cells were washed with FACS buffer and acquired on a BD FACS Canto flow cytometer. Raw data files in FCS format were exported and subsequently analyzed using FlowJo v.10.

### Histopathology

The left lung was immersed in 70% alcohol formalin for 48 hours to facilitate fixation. Subsequently, the tissue was transferred to 10% neutral buffered formalin until further processing for paraffin embedding and sectioning. Sections were stained with hematoxylin and eosin (H&E) and independently evaluated for inflammatory changes by a certified pathologist in a blinded manner. Histopathological images were captured using a LionHeart microscope (BioTek, Agilent).

### Transcriptome analysis using RNA sequencing

RNA was extracted from lung tissue specimens utilizing the RNeasy Plus Mini Kit (Qiagen) according to the manufacturer’s instructions. The quality and integrity of the isolated RNA were assessed using an Agilent Bioanalyzer, and only samples with a RIN value greater than 7 were selected for subsequent library preparation using the NEBNext Ultra II Directional RNA library kit with polyA selection as per the manufacturer’s instructions (NEB, Ipswich, MA). Briefly, two rounds of poly (A) selection were performed on total RNA using paramagnetic oligo (dT) beads. The purified mRNA was fragmented with heat and cations and then converted to cDNA using a mixture of random primers for first-strand synthesis, followed by standard second-strand synthesis. The resulting molecules were ligated to a universal adaptor. The Illumina sequences and the unique index information were added via PCR. The resulting cDNA libraries were quantified using quantitative PCR (qPCR) in a Roche LightCycler 480 with the Kapa Biosystems kit for Illumina library quantification (Kapa Biosystems, Woburn, MA) prior to cluster generation. RNA sequencing was performed on the Illumina NextSeq 500 as described by the manufacturer (Illumina Inc., San Diego, CA).

Raw reads were trimmed usingTrim Galore, (v.0.6.10) and then aligned to the rat reference genome from Ensembl (Rattus norvegicus mRatBN7.2 Release 113) using STAR (v.2.7.11b)^39^. Mapped reads were counted using HTSeq-count (version 2.0.3) followed by normalization, differential gene expression, and pathway analysis using DESeq 2 and IDEP^40–42^.

### Western Blot analyses

Protein from cells or tissue was extracted using a cocktail of RIPA lysis buffer (Thermofisher) and Halt protease and /or phosphatase inhibitor (Thermofisher). Protein concentration was measured using BCA assay (Pierce). For tissue samples, we used a capillary-based protein-resolving simple western technique in a Jess instrument using 12-230kDa fluorescence separation module, 8 x 25 capillary cartridges (ProteinSimple, biotechne) with a protein loading concentration of 1mg/mL and primary mouse anti CFTR antibody (UNC 596, UNC chapel Hill) at dilution of 1:400. For control protein, mouse anti betaactin antibody (Sigma) as primary at a dilution of 1:400 was used. Anti-Mouse secondary antibody was used as secondary antibody along with streptavidin-HRP and Luminol-S substrate (biotechne) for the chemiluminescence-based protein detection. After CFTR detection stripping using Replex buffer (biotechne), the control protein (Beta actin) was detected in the same capillary module. Peak area of the band density was calculated by Compuss for SW software,v.6.1.0 (biotechne) and compared between treatment groups after normalizing with control protein peak area.

### Detection of G551D-CFTR phosphorylation

To understand the effect of Apr on the G551D-CFTR phosphorylation, we performed in vitro experiment using Fisher rat thyroid (FRT) cells expressing G551D-CFTR as described by Wang et al.,^43^ with some modifications. Briefly, we treated cells with either 10uM Apr (in DMSO) or DMSO (vehicle) for 30 min. Cells were washed with ice cold PBS and the protein was extracted using RIPA lysis buffer with Halt protease and phosphatase inhibitor (Thermofisher). After measuring concentration using BCA assay (Pierce), 30 ug protein was used for resolving on Tris-acetate 3-8% gradient gel (Invitrogen) and transferred to PVDF membrane (Immobilon). Total CFTR was probed using rabbit (D6W6L, cell signaling) anti CFTR antibody, and for phosphorylation detection, we used phosphorylation-sensitive monoclonal mouse anti-CFTR antibody UNC 570 ((University of North Carolina at Chapel Hill) at a dilution of 1:5000. Beta-actin was used as a loading control and probed using mouse anti beta actin(sigma) antibody at a dilution of 1:5,000. Goat anti-rabbit IgG (azure spectrum 800, AC2134) and goat polyclonal anti-mouse HRP (Daco) were used as secondary antibodies at a dilution of 1:20,000. For chemiluminescent detection, SuperSignal west Femto chemiluminescent substrate (Thermofisher) was used. Images were acquired using the LiCor Odyssey M system and analyzed using Image Studio.

### Statistics

All the statistical analyses, including the correlation analysis, were performed using GraphPad PRISM v.10. Comparisons between groups were made using either an unpaired Welch’s t-test or the Mann-Whitney test. When the two-tailed p-value was less than or equal to 0.05, we rejected the null hypothesis that the treatment had no effect and considered that the difference between the means of the two groups was statistically significant. For correlation analysis, Pearson’s correlation coefficient (r) was calculated to assess the magnitude and the direction of the correlation. The percentage of variance explained was calculated using the ‘r-squared (r^2^)’ value. A simple linear regression analysis was used to assess goodness of fit with r^2^. A p-value of less than or equal to 0.05 was considered statistically significant.

## Results

### Apremilast preserves lung function in CF rats despite acute P. aeruginosa infection

In this study, we focused on understanding the beneficial properties of inhibiting PDE4 in the management of CF disease. PDE inhibitors have been approved for the treatment of COPD and have been proven to improve lung function and reduce pulmonary exacerbations^44–46^. However, the effect of selective PDE4 inhibition on lung function improvement in CF has not been thoroughly evaluated. To understand the standalone effect of Apr on lung function, we treated CF rats prophylactically before acutely infecting them with *P. aeruginosa* (Fig.1A). Lung function parameters were analyzed using Flexivent oscillometric analysis and compared against sham CF and WT rats.

**Fig. 1.**
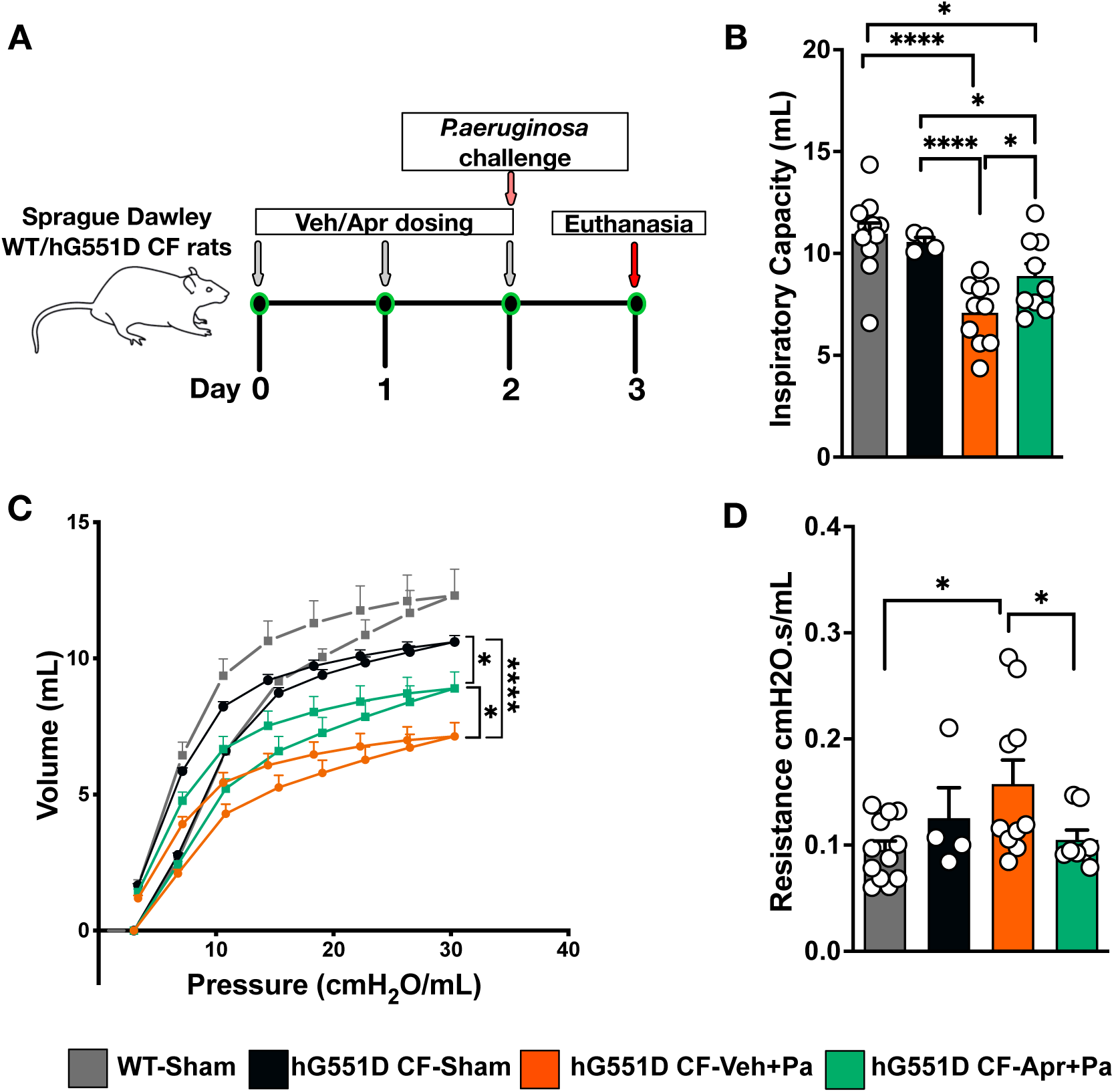
Apremilast (Apr) preserves lung function in cystic fibrosis (CF) rat models following acute *Pseudomonas aeruginosa* (Pa) infection. Selective PDE4 inhibition with a single prophylactic dose of 10 mg/kg per day orally for three days (**A**) attenuates infection-induced increase in airway resistance (**D**), while significantly protecting inspiratory capacity (**B**) and volume at peak pressure in the PV-loop (**C**) as measured by in-vivo respiratory mechanics using Flexivent oscillometry analysis. Data demonstrate the protective effects of Apr on pulmonary physiology under infectious challenge. *p≤0.05, **p≤0.01, ***p≤0.001, ****p≤0.0001.

Analysis of respiratory mechanics 24 hours post-acute *P. aeruginosa* infection in CF rats resulted in a significant reduction of the inspiratory capacity (Fig.1B) and volume at peak pressure in the PV-loop maneuver (Fig.1C) compared to uninfected ones. Irrespective of infection the hysteresis area of CF rats did not show any significant change, but it was significantly reduced compared to WT rats (Fig.1C and Sup. Fig.1A). However, when treated with Apr, these functional parameters remained significantly higher compared to Veh treated CF rats, with a significant reduction in lung resistance as a single compartment, and a substantial decrease in central airway resistance (Fig.1B, C and D, Sup. Fig.1B), indicating the potential role of Apr in protecting lungs from infectious injury that could lead to decreased lung function.

### Apremilast prevents the inflammatory damage caused by acute P. aeruginosa infection in the CF rat lung

PDE inhibitors are known anti-inflammatory agents and have been used in various inflammatory conditions, including chronic respiratory diseases^47–49^. As a selective PDE4 inhibitor, Apr elicits a strong anti-inflammatory effect, which has been utilized for the treatment of chronic inflammatory conditions such as psoriasis and psoriatic arthritis^50^. It offers a wide therapeutic index with fewer side effects than its predecessor drug, roflumilast, approved for COPD^51^. However, studies have shown that immune modulation by roflumilast increased *Pseudomonas* burden in the lungs of WT mice^52^, raising concerns about its use in CF. Nonetheless, the immunoregulatory effect of Apr and its role in bacterial clearance in CF have not yet been studied.

After prophylactic treatment with either Veh or Apr and 24 hours after an acute *P. aeruginosa* infection, the lungs were collected, and histopathological analysis was performed to evaluate the pathological changes. Blinded analysis by an expert pathologist showed a significant decrease in lung inflammation in CF rats that received the Apr treatment (Fig.2A and 2B). However, screening of the BALF did not show any prominent change in the total cell count (Fig.2C) and in cell differentials (Sup. Fig.2A).

**Fig 2.**
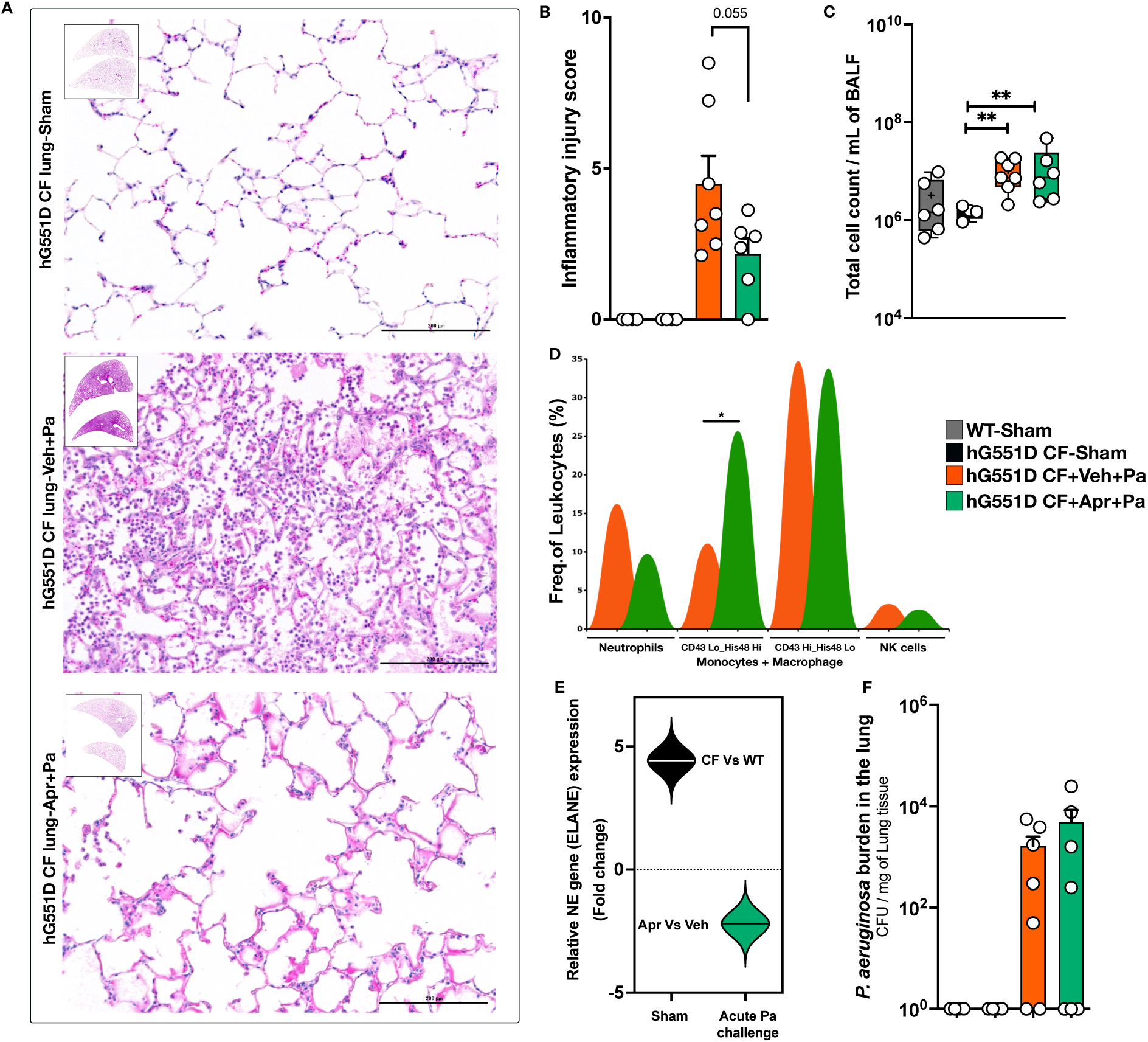
Apremilast (Apr) protects the CF rat lung from inflammatory damage following an acute *P. aeruginosa* (Pa) infection. Hematoxylin and Eosin (H&E) stained lung tissue sections were imaged (**A**) and analyzed by an expert pathologist in a blinded manner to assess the pathological damage caused by the Pa challenge. A decrease in inflammation was observed in the Apr-treated group, as indicated by the pathologist’s score (**B**). The total number of cells present in the BALF was not changed between the Veh and Apr treatments (**C**). However, flow cytometry analysis of the lung tissue showed a decrease in the mean percentage of lung neutrophils and a significant increase in the CD43Lo/His48Hi monocyte/macrophage population (**D**), while differential gene expression analysis showed downregulation of neutrophil elastase in Apr treated CF rats (**E**) but without any significant change in the lung Pa burden (**F**). *p≤0.05, **p≤0.01, ***p≤0.001, ****p≤0.0001.

Since Apr is known for decreasing the infiltration of inflammatory cells^53–55^, we investigated further by performing the flow cytometry on the leukocytes extracted from lung tissue. Although not significant, treatment with Apr tended to decrease the percentage of total neutrophils in the CF lung after acute infection compared to the Veh-treated group. Interestingly, flow cytometry analysis also revealed a significant increase in the percentage of the CD43Lo_His48Hi monocyte/macrophage population (Fig.2D). We therefore interpret neutrophil-directed effects cautiously and integrate them with cytokine and elastase data below.

### Apremilast attenuates inflammation by modulating Th1 and Th17 immune responses but without a significant increase in P. aeruginosa burden

We first defined baseline airway cytokines in sham CF versus WT rats to identify CF-intrinsic inflammation. Analysis of inflammatory cytokines in the BALF of sham CF rats using the ELISA based Luminex assay revealed significant changes in the levels of several cytokines compared to WT controls. A significant increase in IL-6, along with Th2 cytokines IL-4, IL-5, IL-13, and IL-10 was notable, indicating the presence of an altered inflammatory response in the lungs of these preclinical models. On the other hand, IFN-γ and VEGF were decreased in the CF rat BALF. It is intriguing to see that two cytokines, IFN-γ and IP-10, which are closely associated with the Th1 cytokine response, changed in opposite directions (Fig.3 and Sup.Fig.3).

**Fig 3.**
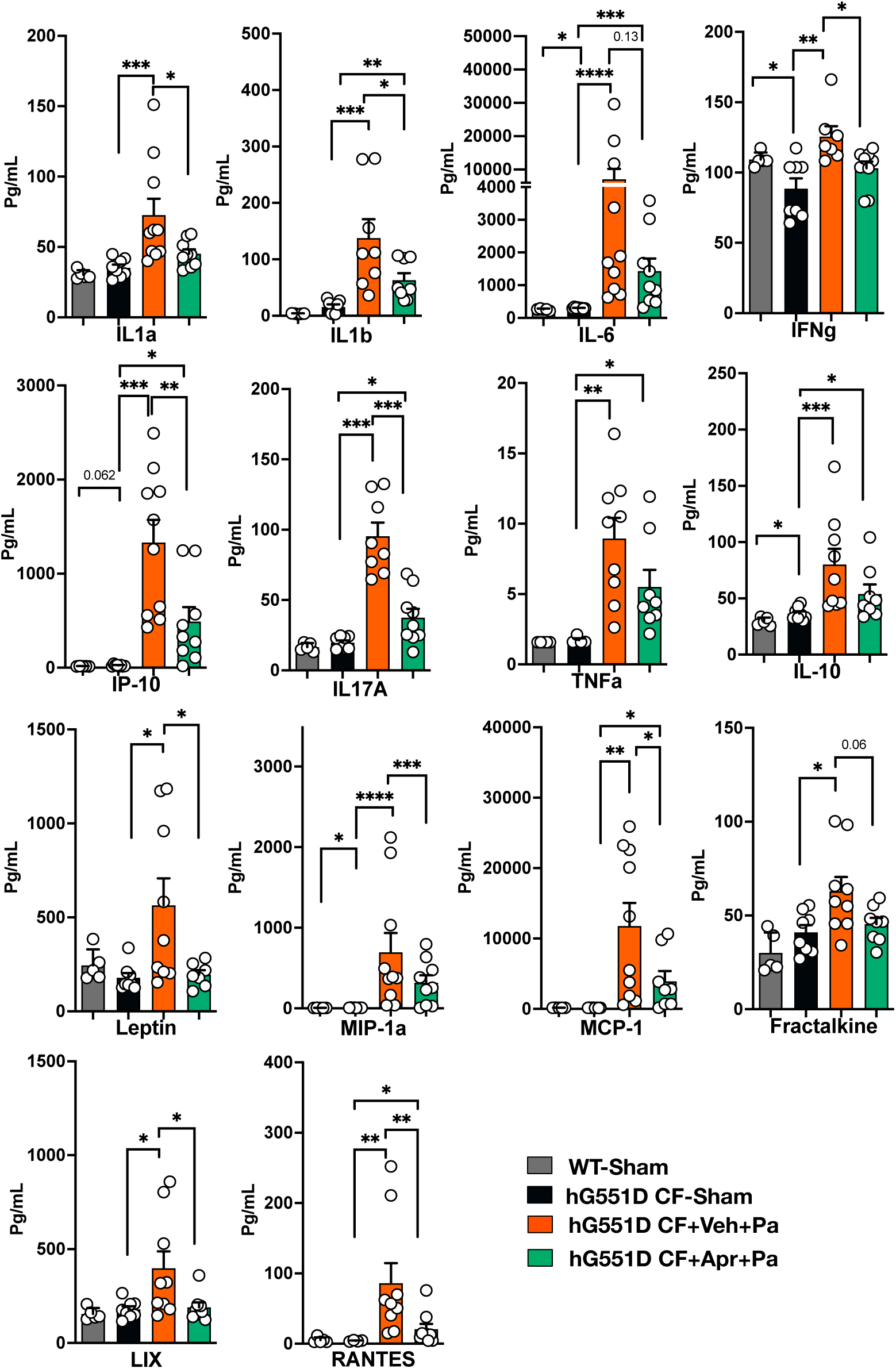
Apremilast (Apr) attenuates CF rat lung injury after an acute *P. aeruginosa* (Pa) infection by modulating inflammatory cytokines. Analysis of cytokine levels in the bronchoalveolar lavage fluid (BALF) using an ELISA based Luminex assay revealed a significant increase in major proinflammatory cytokines following acute infection with Pa. Several of these cytokines were found at significantly reduced levels in CF rats that were prophylactically treated with Apr. *p≤0.05, **p≤0.01, ***p≤0.001, ****p≤0.0001.

We next asked how acute *P. aeruginosa* infection perturbs this milieu in CF and whether Apr prophylaxis rebalances these responses. Hence, we analyzed the cytokine profile in BALF from CF rats treated with and without Apr after acute *P. aeruginosa* infection. While the majority of cytokine levels were increased due to infection, a significant decrease in IL-4 compared to sham CF rats was found noteworthy. Of the 27 inflammatory cytokines analyzed, more than 30% (n=10) showed a significant decrease in the Apr-treated group compared with the veh-treated group (Fig.3). Several of these cytokines, including IL-6 and Fractalkine which showed a decrease after Apr treatment, were associated with either Th1 or Th17 immune responses. Apr is known for inhibiting TNFα ^56^, and its level along with IL-10 decreased in the BALF of Apr treated CF rats, though it was not significant. The Th2 cytokine IL-5 significantly decreased after Apr treatment compared to the Veh group, while IL-4, and IL-13 did not show any significant change (Fig.3 and Sup. Fig.3). Despite this significant reduction of major proinflammatory cytokines in the BALF, the bacterial load in the lung did not show any significant difference between Veh and Apr treated groups (Fig.2F). Thus, Apr rebalanced Th1/Th17- skewed inflammation without compromising pathogen control.

### Differential gene expression analysis confirms regulation of Th1 and Th17 immune responses in CF lung by apremilast

To place these protein-level changes in broader context, we performed RNA-seq on lung tissue from infected CF rats treated with Apr versus Veh. Transcriptomic analysis of the lung tissues reveals that the gene expression profile of the CF rat lung differs from that of WT healthy rat lungs (Fig.4A and B). When there is an acute infection, the profile further changes profoundly, and prophylactic Apr treatment brings the expression profile closer to that of untreated (sham) CF rat lungs (Fig.4B).

**Fig. 4.**
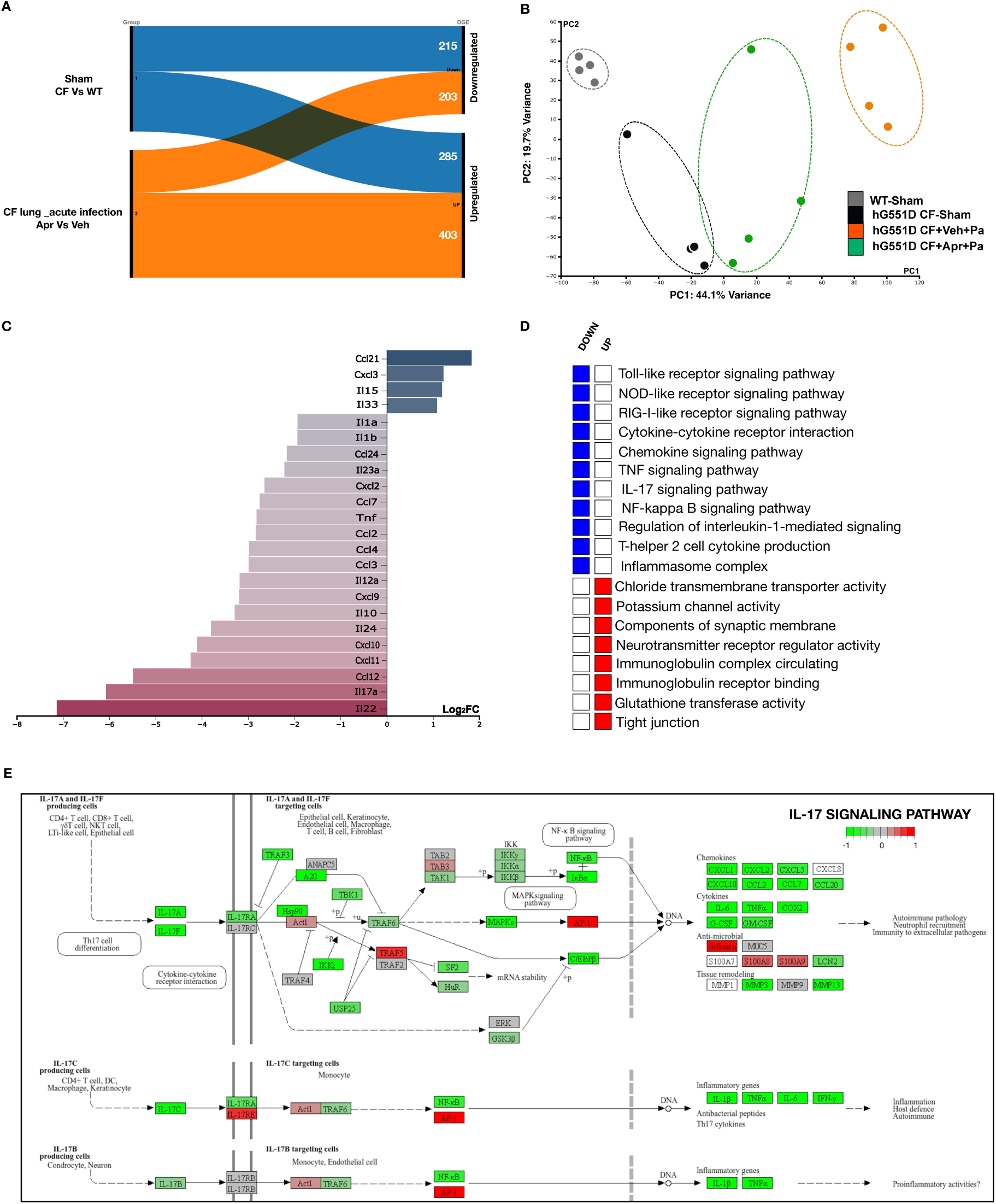
Transcriptomic analysis of the CF rat lung reveals important inflammatory pathways modulated by apremilast (Apr) to reduce the inflammation. Bulk RNA sequencing of lung tissues shows a significant change (>4 FC) in more than 600 genes in the Apr-treated group (A), shifting the gene expression profile more towards the untreated control CF rat lung as shown by the PCA plot (B). The expression of key inflammatory mediators (C) and immune pathways (D) were significantly downregulated by the short-term prophylactic Apr treatment in response to acute *P. aeruginosa* (Pa) infection, of which genes related to the IL-17 signaling pathway are noteworthy as depicted by the KEGG graph rendered by Pathview (E).

Compared to Veh treatment, there were more than 600 differentially expressed genes that showed a fold change (FC) of more than 4, in the Apr-treated group (Fig.4A). Most of the inflammatory mediators that we analyzed using the Luminex assay showed a similar trend in their gene expression pattern based on the transcriptomic analysis (Fig.4C). The expression of IL-1α, IL-1β, IL-17, TNFα, and IL-10 is among those genes that showed significant (p ≤ 0.05) downregulation, further supporting our finding with the Luminex assay that Apr modulates these proinflammatory cytokines to exert its anti-inflammatory effect. Although IL-12a expression was downregulated with Apr, the lack of any significant change in the BALF’s IL- 4 and IL-12p70 protein level (Fig.4C and Sup. Fig.5) led us to emphasize the consistent and significant modulation of Apr within Th17 and IL-1/TNF pathways.

**Figure 5.**
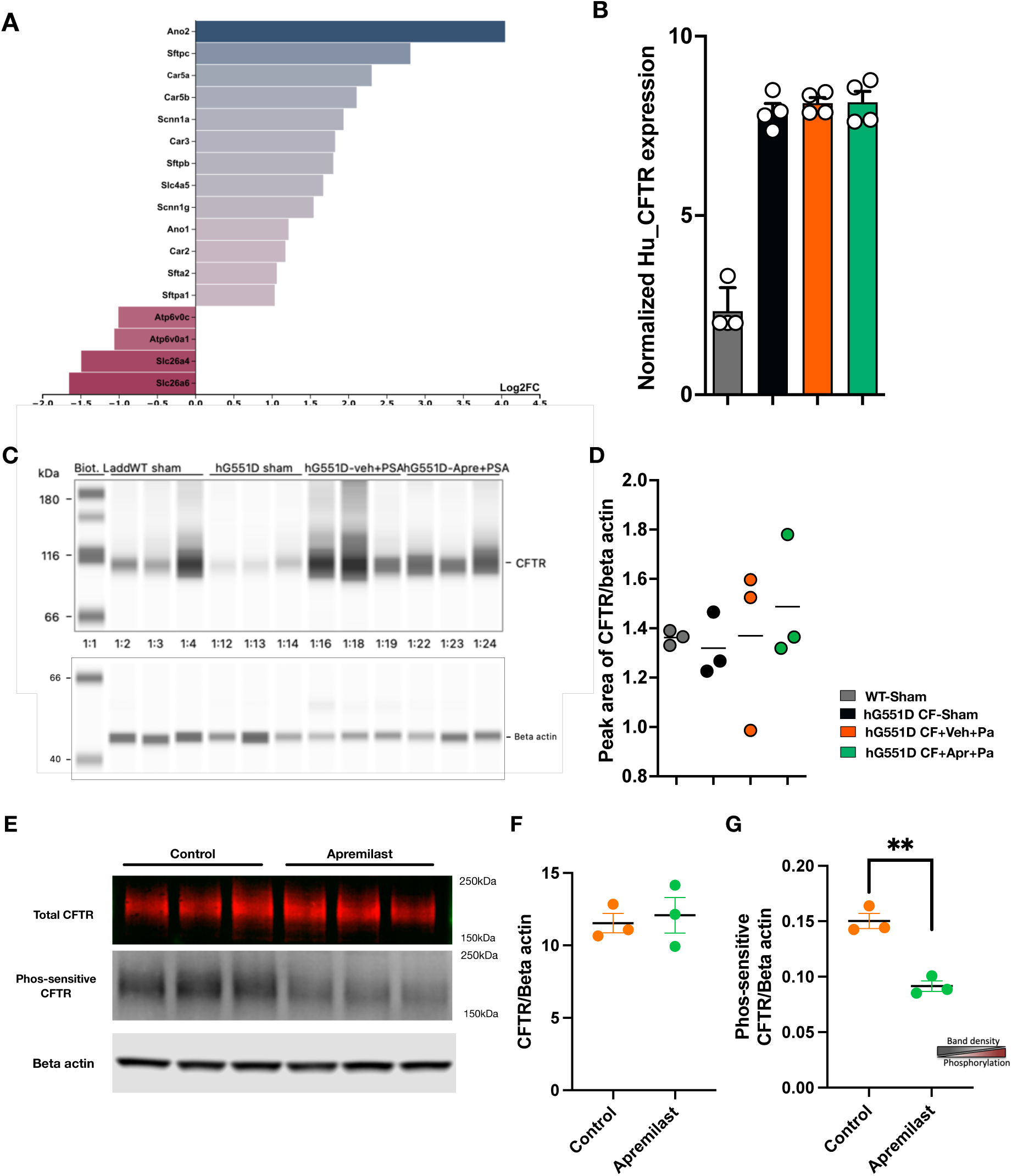
Apremilast protects the CFTR protein in the lungs of CF rats. DGE analysis shows significant changes in the genes related to the pH of the airway surface liquid (ASL) (**A**). Gene expression (**B**) and Western blot (**C** and **D**) analysis of CFTR in the rat lung reveal no significant difference between treatment groups. Western blot analysis of FRT cells expressing human CFTR with the G551D mutation shows significantly increased CFTR phosphorylation after treatment with Apr (10 uM), but no change in total CFTR expression (**E**, **F**, and **G**). *p≤0.05, **p≤0.01, ***p≤0.001, ****p≤0.0001.

Among the significantly changed proinflammatory cytokines, IL-17a and IL-22 were the prominently downregulated ones (Fig.4C). These two cytokines are distinct and belong to separate families, the IL-17 cytokine family and the IL-10 cytokine family respectively. However, they are functionally related and are produced mainly by Th17 cells. A downregulation of these cytokines indicates the suppression of Th17 immune response^57–60^. Pathway analysis shows that several genes in the IL-17 signaling pathway were downregulated when there was Apr treatment (Fig.4E). Notably, IL-17 levels in the BALF also exhibited a weak but significant negative correlation with inspiratory capacity, similar to several other inflammatory cytokines such as IL-1β, IL-13, IL-2, G-CSF, and Eotaxin. However, IL-4, IL-12p70, and VEGF levels showed significant positive correlations (Sup.Fig.4), indicating the critical roles played by the inflammatory milieu in lung function.

### Apremilast shows a broad spectrum of immunomodulatory activity in CF rat lung

Transcriptomics-based pathway analysis showed that Apr exerts its activity from pathogen detection to transcriptional regulation of inflammation. The top 50 pathways identified using the KEGG database showed a significant downregulation of pattern recognition receptors (PRRs) signaling pathways, such as Toll-like receptor (TLR), NOD-like receptor (NLR), RIG-I-like receptor (RLR) signaling pathways, TNF signaling pathway, IL-17 signaling pathway, NF-kappa B signaling pathway, Th1 and Th2 cell differentiation, cytokine-cytokine receptor interaction, and chemokine signaling pathway (Fig.4D and Sup. Fig.5). In summary, Apr could down regulate immune process at pathogen detection, antigen processing and presentation, cellular responses to inter cellular communication via immune mediators. Analysis using other databases (GO) also showed downregulation of cytokine, chemokine, and PRRs activity. Downregulation of NK cell chemotaxis and IL-1-mediated signaling pathway was also noteworthy (sup. Fig.5). Many genes in the IL-17 pathway, one of the key inflammatory mediators and a potential anti-inflammatory therapeutic target in the CF lung^61^, were significantly downregulated after Apr treatment (Fig.4E), indicating a broader modulatory effect on specific inflammatory pathways. A decrease in the number of infiltrated neutrophils (Fig.2D) and downregulation of neutrophil elastase (NE) (Fig.2E) in Apr treated rats could be due to the direct PDE4 inhibitory action in neutrophils, as well as indirectly through regulation of inflammatory cytokine responses.

### Apremilast enhances CF rat airway mucosal defenses through multiple mechanisms

Compared to Veh treatment, a significant upregulation of cytochrome p450 family genes in Apr treated CF rat lung tissue indicates the pharmacological and metabolic activity of this orally administered drug in the lung tissue. A significant upregulation of tight junctions, ion channel activity, especially chloride and potassium ion transporter activity, and ciliary components, suggests that Apr treatment fortifies the mucosal barrier and defense mechanisms. Although we did not observe a significant change in mucin gene expression, we did find a significant change in surfactant protein gene expression (Sftpa1, Sftpb, Sftpc, and Sfta2) (Fig.5A).

Since the pH of the ASL plays a critical role, we compared the expression of genes important for maintaining the pH of ASL, between the Veh and Apr treatment groups after an acute *P. aeruginosa* infection. Interestingly, we found a significant decrease in the expression of genes Atp6v0c and Atp6v0a1(Fig.5A), which encode components of vacuolar H^+^-ATPase (V-ATPase), a major H^+^-secreting mechanism in the lower airways^62, 63^. Since the pH of ASL is determined by the acid (H+) and base (HCO3^-^) secretion^63, 64^, we looked at important transporters of HCO3- at the apical surface in the lower airways. Intriguingly, we found an upregulation of the genes Ano1 and Ano2, which encode for Ca-activated chloride channels (CaCC), one of the major HCO3- transporters in the small airways. CFTR forms another important HCO3-transporter in the apical side^63, 65^; but there was no significant change in its expression based on the transcriptomic data (Fig.5B). However, we found a significant increase in the expression of Slc4a5 (p adj=0.019), which encodes a member of the solute carrier family 4 proteins. The transporters of this family are responsible for the Na+-HCO3^-^cotransport (NBC) activity, where a gradient of Na+ is generated for the co-transport of HCO3- into the cell at the basolateral surface and thereby supplies HCO3- to apical transporters. We believe that an upregulation of sodium channel epithelial 1 subunit alpha (Scnn1a) and gamma (Scnn1g) (Fig.5A), which are subunits of ENac channels responsible for the Na+ transport into cells, could be part of creating a Na+ gradient to regulate the alveolar fluid clearance (AFC). An upregulation of carbonic anhydrase (CA) genes (Car2, Car3, Car5a, and Car5b) indicates endogenous production of bicarbonate for its secretion to the apical transporters as a part of alkalinizing the ASL(Fig.5A). On the contrary, Slc26 family transporters (Slc26a4 and Slc26a6), which are important for non-CFTR mediated HCO3- transport, were found to be significantly downregulated in the Apr-treated group compared to the Veh group (Fig.5A).

### Modulation of inflammation did not affect the CFTR expression in the CF rat lung

Cellular CFTR expression has been shown to be influenced by inflammatory cytokines. Earlier studies showed that cytokines, like IFN-γ and TNF-α, decreased CFTR expression when cell cultures were treated with those cytokines individually^66, 67^. Conversely, recent research indicates that a combination of certain cytokines, such as TNF-α and IL-17, upregulates CFTR expression and alkalinizes the ASL^63, 68^. While these in vitro findings highlight the impact of inflammation on CFTR expression, it remains unclear how specific pathogen infection and the associated complex inflammatory responses mediated by several cytokines affect the CFTR expression in CF lung tissue. More importantly, it is critical to see whether attenuation of inflammation mediated by Apr has any unintended effects on the CFTR expression. Interestingly, our RNA sequencing based gene expression analysis and western blot based protein expression analysis did not show any significant change in the expression of CFTR in the CF rat lung after an acute *P. aeruginosa* infection with or without Apr prophylactic treatment (Fig.5B, C, and D). Despite a broad modulation of the inflammatory responses, particularly significant downregulation of several proinflammatory cytokines, including TNF-α and IL-17, Apr treatment did not cause any decrease in the CFTR expression both at the mRNA and protein levels.

### Apremilast increases CFTR phosphorylation in cells with G551D mutation

The G551D mutation is a class III CFTR mutation known to impair PKA dependent activation of CFTR due to impaired channel phosphorylation^43^. Given that Apr inhibits PDE4, and the resulting increase in cAMP leads to activation of CFTR via PKA, the mechanism by which Apr activates CFTR in the context of the G551D mutation needs investigation. Hence, we validated the effect of Apr on phosphorylation of CFTR using the same in-vitro model used by wang *et al* to assess the impact of the G551D mutation on CFTR activation^43^. When the western blot was performed using the same phosphorylation sensitive antibody probe (UNC 570, epitope: aa731–742), treatment with Apr at 10uM for 30 minutes significantly increased the phosphorylation of CFTR in FRT cells carrying human CFTR gene with G551D mutation (Fig.5E, F, and G). This change in phosphorylation is identified by a significant decrease in the band density as this probe cannot bind its target when there is phosphorylation at ser 737. The result shows greater implication of Apr combination therapy with CFTR modulators such as Iva, since channel phosphorylation has an impact on the stimulation rate of Iva.

## Discussion

Due to adverse side effects and incompatibility with CFTR modulators, approved anti-inflammatory therapeutic options in CF care are very limited. Considering this, we focus on testing the potential of anti-inflammatory drugs that do not belong to the steroidal or non-steroidal classes but have been approved for other chronic inflammatory conditions leveraging the benefits of the drug repurposing strategy^69^.

Selective phosphodiesterase (PDE) inhibitors have fewer side effects than non-specific inhibitors and have been approved for several chronic inflammatory conditions. Out of 11 isoforms, PDE4 is the most predominantly expressed isoform in lung epithelial cells^70^. Hence, selective inhibition of PDE4 has been approved for respiratory diseases, such as COPD (e.g., Roflumilast) and IPF (e.g., Nerandomilast). Recently, drugs that can selectively inhibit multiple PDE isoforms have been approved for COPD (e.g., Ensifentrine, a PDE3 and 4 inhibitor). Although respiratory disease manifestation contributes to morbidity and mortality in CF, its multiorgan involvement and compromised immune defense system raise concerns about the use of these small-molecule anti-inflammatory drugs in CF. Reports from preclinical studies, such as an increase in *Pseudomonas* burden in WT mice lungs after treatment with a more potent roflumilast^52^, support this concern and keep all members of this group on the risky margin. However, we believe that because of several unknown mechanisms of action of the parent compound and its metabolites, other than inhibition of PDE4 and the associated increase in the cellular cAMP, the therapeutic effect of these drug molecules might vary.

Apremilast (Apr) is a selective PDE4 inhibitor approved for psoriasis and psoriatic arthritis for patients ≥6 years old a decade ago^54, 71^. Although less potent than roflumilast, it showed a wider therapeutic index than its side effects and produced effects similar to other PDE4 inhibitors, especially in reducing neutrophilic pulmonary inflammation in preclinical models^51^. Proinflammatory cytokines such as IL-1, IL-6, IL-17, and TNF-α are key mediators of inflammation in CF. Apr is a known modulator of IL-17 and TNF-α, as well as other cytokines^54, 55, 72^. However, the efficacy of Apr in modulating inflammatory mediators hasn’t been tested under CF conditions, particularly in an infection scenario. Hence, we evaluated the immunomodulatory properties of Apr when administered as a short-term prophylactic dose to CF rats before an acute infection with *P. aeruginosa* and its effects on lung function and infectious burden.

Apr modulated immune responses across multiple nodes without increasing bacterial burden (Figs. 2–4). In the CF lung, prophylactic Apr curtailed the infection-induced surge of IL-17A, a Th17 cytokine central to neutrophil recruitment and antibacterial defense. Because excessive IL-17A promotes tissue injury, a balanced IL-17A response is critical^73–75^. Significantly elevated IL-17A, both at the mRNA and protein levels, was demonstrated in the sputum of CF patients, and it was significantly higher when there was chronic colonization of *Pseudomonas* compared to never or intermittent colonizers^76^. Corroboratively, our CF rat infection model recapitulates this phenomenon, with a significant increase in IL-17A at both the gene expression level in lung tissue and the protein level in the BALF (Fig.3). Challenge with *P. aeruginosa* caused more than a 380% increase (5 times) in IL-17A in the BALF compared to sham CF rats. Prophylactic treatment with Apr rebalances IL-17A response by downregulating its gene expression and decreasing its protein level by more than 60% compared to the Veh group, yet it remains nearly 90% higher than sham controls. Deficiency of IL-17 leads to increased susceptibility to infections, including *Pseudomonas,* and to an increased pathogen burden, which worsens disease severity^77, 78^. Nonetheless, the reduction of IL-17 in the CF rat lung did not cause any significant change in the lung bacterial burden (Fig.2F), indicating that the attenuated level of IL-17A is still adequate to limit the pathogen burden.

In CF lung disease, IL-17A, produced by both innate and adaptive immune cells contributes to inflammatory damage and has been suggested as a target for anti-inflammatory therapy^79^. In vivo neutralization of IL17A has been shown to promote the resolution of inflammation in a bleomycin induced acute inflammation model^80^. Hence, Apr mediated reduction in IL17A response might be a reason for attenuated lung pathology in Apr treated CF rats. As shown in other preclinical models^51, 81^, Apr downregulated neutrophil elastase in lung tissue and trended toward fewer tissue neutrophils, alongside decreased IL-17A and G-CSF (Fig.2, Fig.3). While flowcytometry-based neutrophil reductions were not significant, the collective data support dampened neutrophil-driven inflammation. We believe that the reduction in neutrophil infiltration could be partly due to decreased IL-17 and its effect on the release of neutrophil chemokines, such as IL-6 and G-CSF (Fig.3, 4C and Sup.Fig.3)^82^. Together with IL-22, IL-17 forms the signature cytokines of type 3 cell mediated effector immunity against extracellular bacteria^83^. Interestingly, IL-22 expression was significantly downregulated along with IL-17 after Apr treatment (Fig.4C), indicating their common cellular origin and complementary functional roles in CF lung’s responses to infection.

In pwCF, respiratory infections markedly accelerate the disease progression and deteriorate the lung function^23^. Our CF rat acute infection model mirrors this scenario and without infection the inflammatory responses in the CF lung, compared to WT rat lungs, were moderate and may not fully capture the true effect of Apr. Although, the improvement of lung function by PDE4 inhibitors is well documented in other respiratory diseases like COPD^44, 84^, it has not been thoroughly tested in CF, especially with Apr. When we treated the rats with short term Apr, it significantly prevented the loss of lung function compared to Veh group after *Pseudomonas* challenge (Fig.1). The levels of several inflammatory mediators including IL-17 in the BALF showed a weak but significant negative correlation with lung functions (Sup.Fig.4) implying that alleviating inflammation and the associated release of mediators by Apr might be benefitted to prevent lung function deterioration.

Pulmonary inflammation plays a critical role in the alveolar surface liquid (ASL) homeostasis. Inflammatory mediators affect ion channels on the apical and basolateral surfaces of epithelial cells and influence fluid transport^85^. For example, IL-1β and IL-4 decrease epithelial sodium channel (ENaC) expression during bacterial infection^86, 87^. The activity of ENaC is important in alveolar fluid clearance (AFC) to regulate the pulmonary edema caused by infection mediated acute lung injury (ALI)^85, 88^. Cyclic AMP increases ENaC activity indirectly and increases the highly selective cation channel density on the cell surface as well as channel open probability of non-selective cationic channels^89–91^. This might explain the reason for a significantly upregulated gene expression of ENaC channel subunits alpha (Scnn1a) and gamma (Scnn1g) (Fig.5). Upregulation of Scnn1a and Scnn1g with Apr (Fig. 5) may therefore support AFC in the infected lung. However, in CF airways, excessive ENaC activity is typically pathologic for mucus hydration on the airway surface^92^. Our transcriptomics data do not resolve epithelial compartments; functional studies are needed to determine net effects on airway hydration versus AFC.

The pH of the ASL is a key factor in host defense against respiratory infections. Inflammatory mediators play an important role in maintaining the pH of ASL; however, studies show that they have pleiotropic effects. For example, while the combination of IL-17A and TNF-α alkalinizes the ASL via CFTR and pendrin, it also upregulates the proton pump ATP12A on the apical surface of the airway epithelial cells^68, 92^. The same combination of cytokines influenced the expression of CFTR and epithelial cell response to CFTR modulators^36, 93^, indicating the impact of immune modulation on ion-channel activity and its implications for host defense mechanisms in CF airways extend beyond just reducing inflammation. Despite a decrease in the expression of IL-17, TNF-α and several other cytokines, Apr treatment significantly upregulated several anionic channels and downregulated proton pumps, but without diminishing CFTR expression at both gene and protein expression levels, suggesting Apr’s potential to enhance the mucosal defense while attenuating the inflammatory responses.

In summary, Apr preserved lung function and mitigated lung injury in G551D CF rats acutely challenged with *P. aeruginosa*, broadly rebalancing Th17/Th1-skewed inflammation without increasing bacterial burden. Apr did not reduce CFTR expression in vivo and increased G551D-CFTR phosphorylation in vitro, highlighting mechanistic compatibility with CFTR modulators. Limitations include the acute infection and prophylactic dosing paradigm, lack of direct ASL pH or mucociliary functional measurements, and transcript-level inference for epithelial transport. Future studies should test therapeutic (post-infection) dosing, chronic and mucoid *P. aeruginosa* models, and combination with ETI to determine clinical translatability.

## Acknowledgements

This work was supported in part by UAB Cystic Fibrosis Center grant (P30DK072482), UAB Flow Cytometry Service Core (AI027767, CA013148, S10OD032296), grants (NIH R01AA027528, CFF HARRIS23G0) to S. V. Raju, and CFF RDP training award (DAVIS24RO), and CFRI Patrick Nash Fellowship to L. Antony. The authors thank Drs. James M. Donahue and Pornima Phatak (VAHCS, Birmingham) for their generous support and assistance with using the Jess (ProteinSimple) instrument. Sponsors had no role in study design, data interpretation, and publication.

