## Supplementary data for "PDE4 Inhibitor Apremilast Rebalances Inflammatory Responses to Pseudomonas aeruginosa Infection in CF Rats"

#### Supplementary figure legends

**Sup. Fig.1.** Flexivent oscillometry-based lung function analysis showed a decrease in hysteresis area of the PV loop after an acute infection with *P. aeruginosa* (Pa) in CF rats, while it was prevented in rats treated with Apr (**A**). Although not significant compared to sham control CF rats, acute infection with Pa increased the central airway resistance. However, prophylactic treatment with Apr prevented this increase in airway resistance (**B**). Data demonstrate the protective effects of Apremilast on pulmonary physiology under infectious challenge. *$p \leq 0.05$, **$p \leq 0.01$, ***$p \leq 0.001$, ****$p \leq 0.0001$.

**Sup. Fig.2**. Differential cell count analysis of the BALF cells, after fixing on the slide using cytospin and staining, showed no significant change in the average percentage of the various cell types.

**Sup. Fig. 3.** Changes in the level of inflammatory cytokines in the bronchoalveolar lavage fluid (BALF) collected from the CF rat lung**.** Prophylactic treatment with Apr for a short term modulated the inflammatory response in CF rat lungs after an acute infection with Pa, as reflected in changes in BALF levels of inflammatory cytokines. *$p \leq 0.05$, **$p \leq 0.01$, ***$p \leq 0.001$, ****$p \leq 0.0001$.

**Sup. Fig. 4.** Correlation of lung functions with cytokine levels in the BALF. Measurement of Pearson's correlation coefficient (R) and simple linear regression analysis on samples acutely infected with *P. aeruginosa* (Pa) revealed a significant correlation of the inspiratory capacity of the CF rat lung with levels of some of the inflammatory cytokines in the BALF. Analysis was performed, including both veh and Apr treatment group samples. A p-value of less than or equal to 0.05 is considered significant ($p \leq 0.05$).

**Sup. Fig.5**. Major pathways that were significantly changed in Apr-treated groups compared to the Veh group, identified using different databases, and are mentioned as abbreviations. The size of the node indicates the number of genes in each pathway. Color indicates the direction of gene expression. The full terms of the abbreviations are provided in the Sup. Table.

#### Supplementary Figure 1

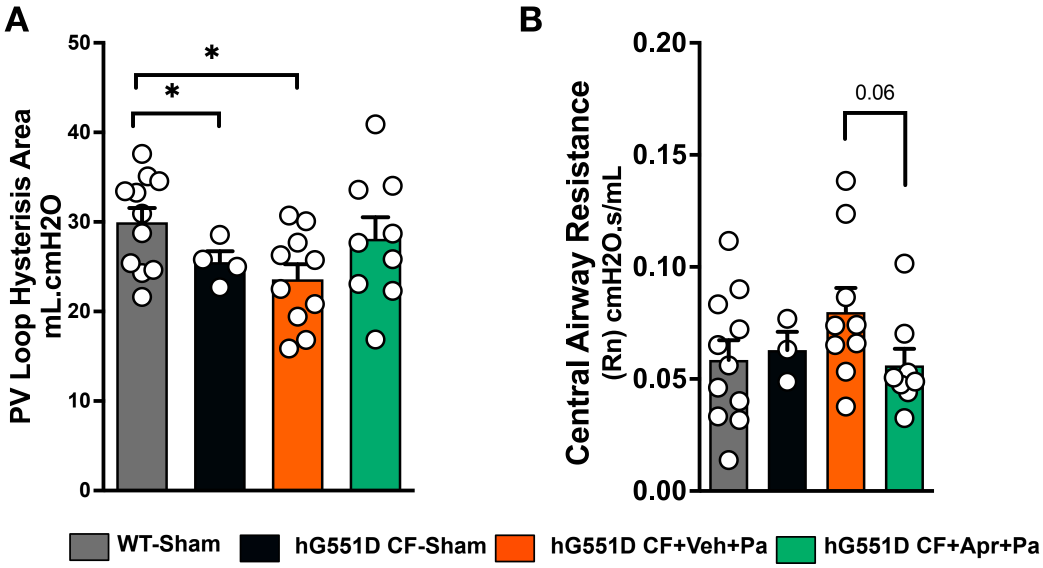

#### Supplementary Figure 2

**A**

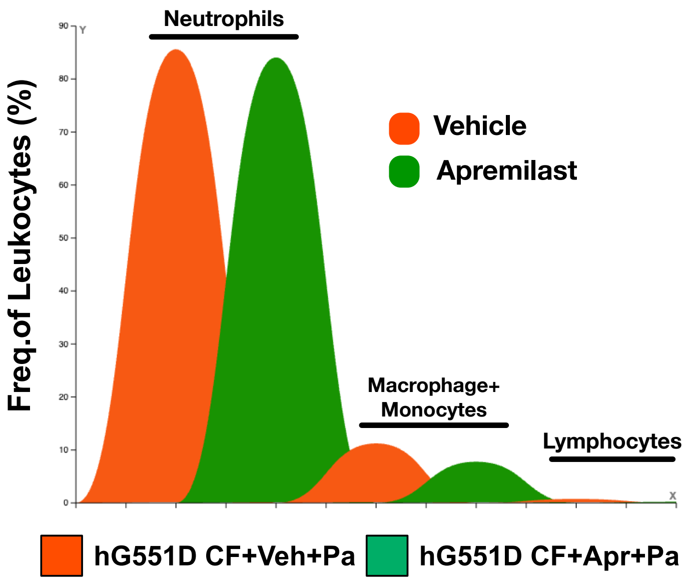

#### Supplementary Figure 3

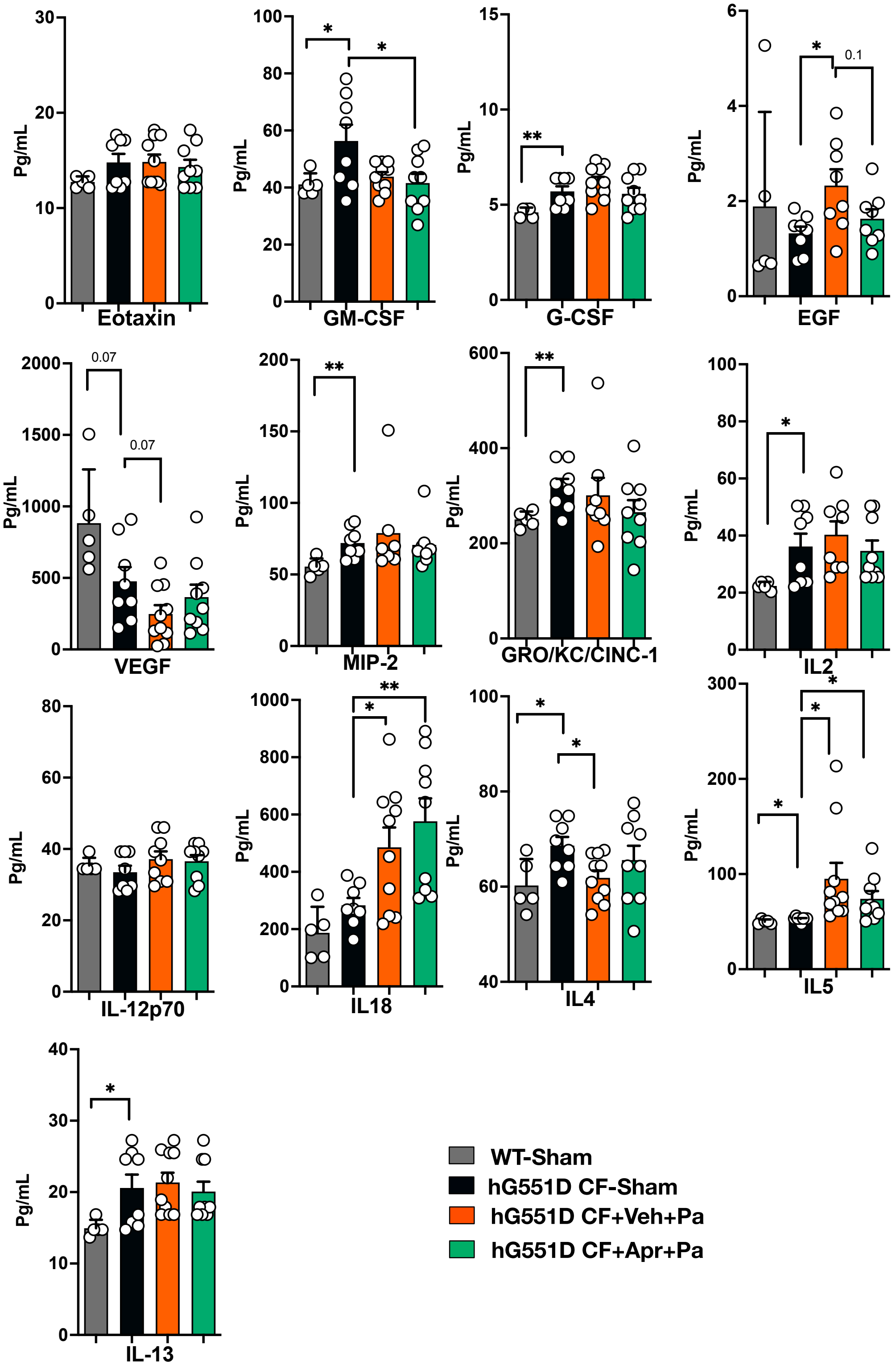

#### Supplementary Figure 4

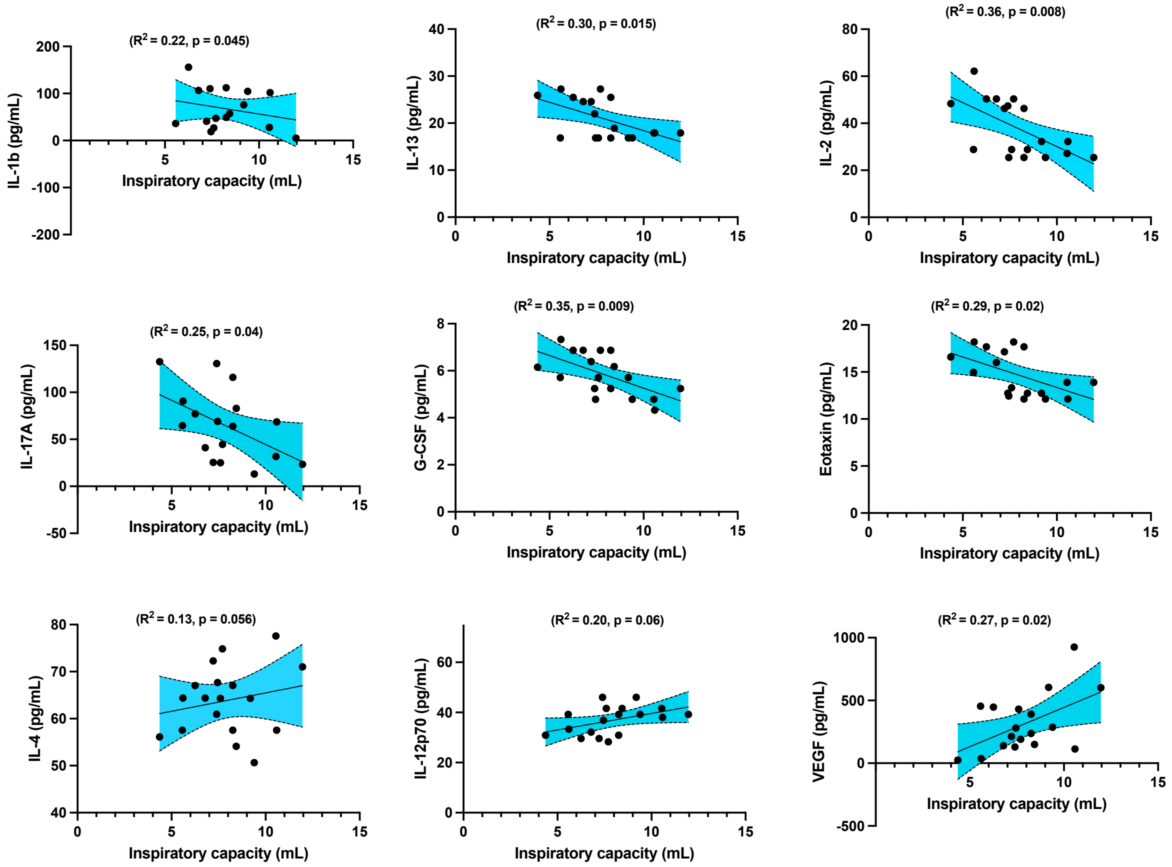

### Supplementary Figure 5

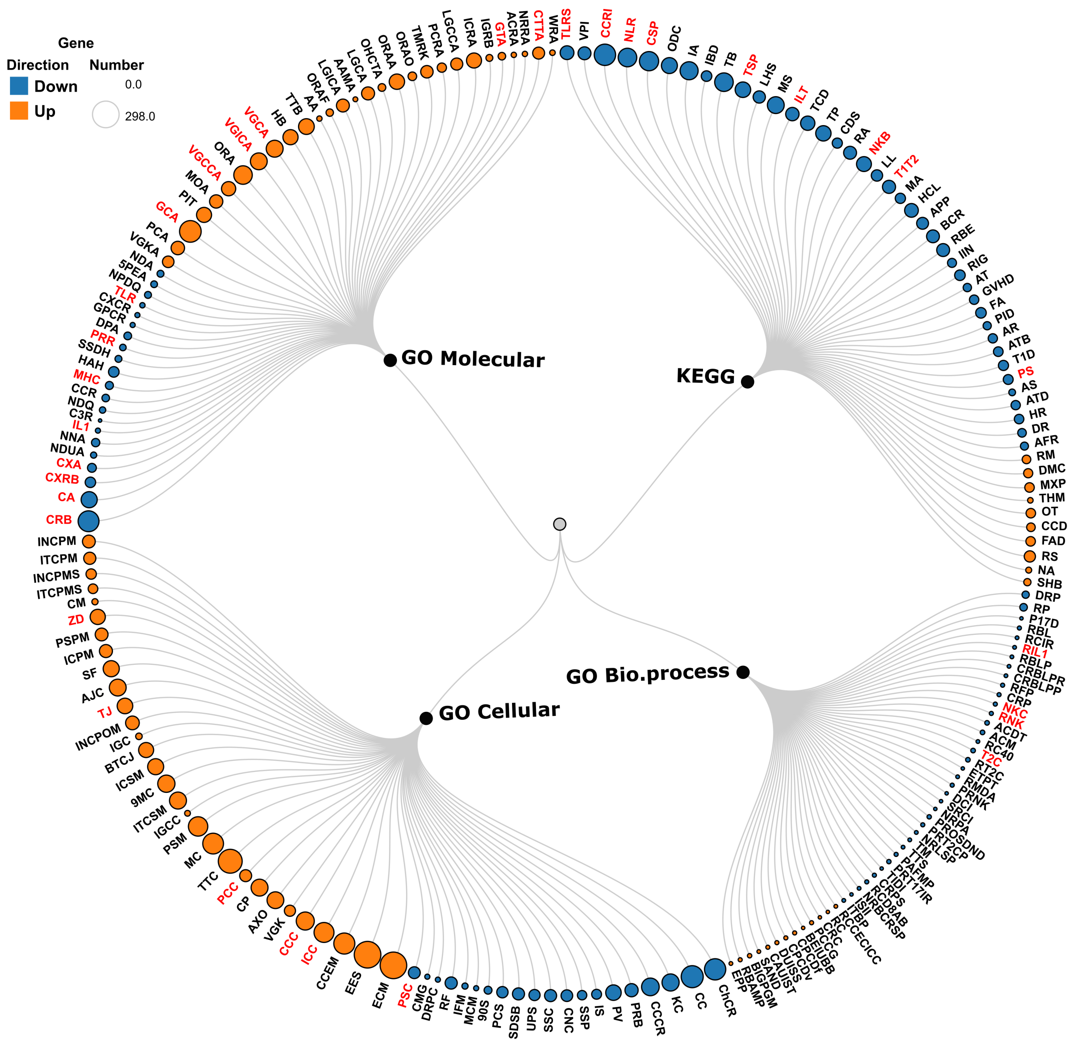

**Supplementary Table**
Details of pathways shown in Sup. Fig. 5

| | |
|---|---|
| **Pathway Name** | **Abbreviation** |
| Toll-like receptor signaling pathway | TLRS |
| Viral protein interaction with cytokine and cytokine receptor | VPI |
| Cytokine-cytokine receptor interaction | CCRI |
| NOD-like receptor signaling pathway | NLR |
| Chemokine signaling pathway | CSP |
| Osteoclast differentiation | ODC |
| Influenza A | IA |
| Inflammatory bowel disease | IBD |
| Tuberculosis | TB |
| TNF signaling pathway | TSP |
| Leishmaniasis | LHS |
| Measles | MS |
| IL-17 signaling pathway | ILT |
| Th17 cell differentiation | TCD |
| Toxoplasmosis | TP |
| Cytosolic DNA-sensing pathway | CDS |
| Rheumatoid arthritis | RA |
| NF-kappa B signaling pathway | NKB |
| Legionellosis | LL |
| Th1 and Th2 cell differentiation | T1T2 |
| Malaria | MA |
| Hematopoietic cell lineage | HCL |
| Antigen processing and presentation | APP |
| B cell receptor signaling pathway | BCR |
| Ribosome biogenesis in eukaryotes | RBE |
| Intestinal immune network for IgA production | IIN |
| RIG-I-like receptor signaling pathway | RIG |
| African trypanosomiasis | AT |
| Graft-versus-host disease | GVHD |
| Fanconi anemia pathway | FA |
| Primary immunodeficiency | PID |
| Allograft rejection | AR |
| Aminoacyl-tRNA biosynthesis | ATB |
| Type I diabetes mellitus | T1D |

| | |
|---|---|
| Proteasome | PS |
| Asthma | AS |
| Autoimmune thyroid disease | ATD |
| Homologous recombination | HR |
| DNA replication | DR |
| Antifolate resistance | AFR |
| Retinol metabolism | RM |
| Drug metabolism-cytochrome P450 | DMC |
| Metabolism of xenobiotics by cytochrome P450 | MXP |
| Taurine and hypotaurine metabolism | THM |
| Olfactory transduction | OT |
| Chemical carcinogenesis-DNA adducts | CCD |
| Fatty acid degradation | FAD |
| Renin secretion | RS |
| Nicotine addiction | NA |
| Steroid hormone biosynthesis | SHB |
| Defense response to protozoan | DRP |
| Response to protozoan | RP |
| Positive regulation of T-helper 17 cell differentiation | P17D |
| Response to bacterial lipoprotein | RBL |
| Regulation of chronic inflammatory response | RCIR |
| Regulation of interleukin-1-mediated signaling pathway | RIL1 |
| Response to bacterial lipopeptide | RBLP |
| Cellular response to bacterial lipoprotein | CRBLPR |
| Cellular response to bacterial lipopeptide | CRBLPP |
| Replication fork protection | RFP |
| Cellular response to peptidoglycan | CRP |
| Natural killer cell chemotaxis | NKC |
| Regulation of natural killer cell chemotaxis | RNK |
| Activation-induced cell death of T cells | ACDT |
| Astrocyte cell migration | ACM |
| Regulation of CD40 signaling pathway | RC40 |
| T-helper 2 cell cytokine production | T2C |
| Regulation of T-helper 2 cell cytokine production | RT2C |
| Electron transport coupled proton transport | ETPT |
| Regulation of myeloid dendritic cell activation | RMDA |
| Positive regulation of natural killer cell mediated cytotoxicity | PRNK |
| Detoxification of copper ion | DCI |
| Stress response to copper ion | SRCI |

| | |
|---|---|
| Negative regulation of plasminogen activation | NRPA |
| Positive regulation of oxidative stress-induced neuron death | PROSDND |
| Positive regulation of T-helper 2 cell cytokine production | PRT2CP |
| Negative regulation of lipopolysaccharide-mediated signaling pathway | NRLSP |
| Thymocyte migration | TM |
| TRIF-dependent toll-like receptor signaling pathway | TTS |
| Platelet activating factor metabolic process | PAFMP |
| Positive regulation of T-helper 17 type immune response | PRT17IR |
| Tolerance induction dependent upon immune response | TIDI |
| Cellular response to progesterone stimulus | CRPS |
| Regulation of CD8-positive alpha-beta T cell differentiation | RCD8AB |
| Negative regulation of B cell receptor signaling pathway | NRBCRSP |
| Intracellular sequestering of iron ion | ISII |
| Inositol trisphosphate biosynthetic process | ITBP |
| Regulation of cell communication by electrical coupling involved in cardiac conduction | RCCECICC |
| Response to cGMP | RC |
| Cellular response to cGMP | CRC |
| Peptide cross-linking via chondroitin 4-sulfate glycosaminoglycan | PCCG |
| Branch elongation involved in ureteric bud branching | BEIUBB |
| Cardiac pacemaker cell differentiation | CPCDf |
| Cardiac pacemaker cell development | CPCDv |
| Dopamine uptake involved in synaptic transmission | DUISS |
| Catecholamine uptake involved in synaptic transmission | CAUIST |
| Sinoatrial node development | SAND |
| Branching involved in prostate gland morphogenesis | BIGPGM |
| Regulation of bile acid metabolic process | RBAMP |
| Epoxygenase P450 pathway | EPP |
| Chromosome centromeric region | ChCR |
| Condensed chromosome | CC |
| Kinetochore | KC |
| Condensed chromosome centromeric region | CCCR |
| Preribosome | PRB |
| Phagocytic vesicle | PV |
| Immunological synapse | IS |
| Small-subunit processome | SSP |
| Condensed nuclear chromosome | CNC |
| Spliceosomal snRNP complex | SSC |
| U2-type precatalytic spliceosome | UPS |

| | |
|---|---|
| Site of double-strand break | SDSB |
| Precatalytic spliceosome | PCS |
| 90S preribosome | 90S |
| MCM complex | MCM |
| Inflammasome complex | IFM |
| Replication fork | RF |
| DNA replication preinitiation complex | DRPC |
| CMG complex | CMG |
| Proteasome complex | PSC |
| Extracellular matrix | ECM |
| External encapsulating structure | EES |
| Collagen-containing extracellular matrix | CCEM |
| Ion channel complex | ICC |
| Cation channel complex | CCC |
| Voltage-gated potassium channel complex | VGK |
| Axoneme | AXO |
| Ciliary plasm | CP |
| Potassium channel complex | PCC |
| Transmembrane transporter complex | TTC |
| Motile cilium | MC |
| Postsynaptic membrane | PSM |
| Immunoglobulin complex circulating | IGCC |
| Intrinsic component of synaptic membrane | ITCSM |
| 9+2 motile cilium | 9MC |
| Integral component of synaptic membrane | ICSM |
| Bicellular tight junction | BTCJ |
| Immunoglobulin complex | IGC |
| Intrinsic component of postsynaptic membrane | INCPOM |
| Tight junction | TJ |
| Apical junction complex | AJC |
| Sperm flagellum | SF |
| Integral component of postsynaptic membrane | ICPM |
| Postsynaptic specialization membrane | PSPM |
| Z disc | ZD |
| Compact myelin | CM |
| Integral component of postsynaptic specialization membrane | ITCPMS |
| Intrinsic component of postsynaptic specialization membrane | INCPMS |
| Integral component of presynaptic membrane | ITCPM |
| Intrinsic component of presynaptic membrane | INCPM |

| | |
|---|---|
| Cytokine receptor binding | CRB |
| Cytokine activity | CA |
| Chemokine receptor binding | CXRB |
| Chemokine activity | CXA |
| NADH dehydrogenase (ubiquinone) activity | NDUA |
| NAD+ nucleosidase activity | NNA |
| Interleukin-1 receptor binding | IL1 |
| CXCR3 chemokine receptor binding | C3R |
| NADH dehydrogenase (quinone) activity | NDQ |
| CCR chemokine receptor binding | CCR |
| MHC protein binding | MHC |
| Hydrolase activity hydrolyzing N-glycosyl compounds | HAH |
| Single-stranded DNA helicase activity | SSDH |
| Pattern recognition receptor activity | PRR |
| DNA polymerase activity | DPA |
| G protein-coupled purinergic nucleotide receptor activity | GPCR |
| CXCR chemokine receptor binding | CXCR |
| Toll-like receptor binding | TLR |
| NAD(P)H dehydrogenase (quinone) activity | NPDQ |
| 5-prime-3-prime exonuclease activity | 5PEA |
| NADH dehydrogenase activity | NDA |
| Voltage-gated potassium channel activity | VGKA |
| Potassium channel activity | PCA |
| Gated channel activity | GCA |
| Potassium ion transmembrane transporter activity | PIT |
| Monooxygenase activity | MOA |
| Voltage-gated cation channel activity | VGCCA |
| Oxidoreductase activity acting on paired donors with incorporation or reduction of molecular oxygen | ORA |
| Voltage-gated ion channel activity | VGICA |
| Voltage-gated channel activity | VGCA |
| Heme binding | HB |
| Tetrapyrrole binding | TTB |
| Aromatase activity | AA |
| Oxidoreductase activity acting on paired donors with incorporation or reduction of molecular oxygen reduced flavin or flavoprotein as one donor and incorporation of one atom of oxygen | ORAF |
| Ligand-gated ion channel activity | LGICA |
| Arachidonic acid monooxygenase activity | AAMA |

| | |
|---|---|
| Ligand-gated channel activity | LGCA |
| Organic hydroxy compound transmembrane transporter activity | OHCTA |
| Oxidoreductase activity acting on the CH-OH group of donors NAD or NADP as acceptor | ORAA |
| Oxidoreductase activity acting on paired donors with incorporation or reduction of molecular oxygen NAD(P)H as one donor and incorporation of one atom of oxygen | ORAO |
| Transmembrane receptor protein kinase activity | TMRK |
| Potassium channel regulator activity | PCRA |
| Ligand-gated cation channel activity | LGCCA |
| Ion channel regulator activity | ICRA |
| Immunoglobulin receptor binding | IGRB |
| Glutathione transferase activity | GTA |
| Acetylcholine receptor regulator activity | ACRA |
| Neurotransmitter receptor regulator activity | NRRA |
| Chloride transmembrane transporter activity | CTTA |
| Wnt-activated receptor activity | WRA |
